# Removal of NHS-labelling By-products in Proteomic Samples

**DOI:** 10.1101/2024.08.15.607975

**Authors:** Yana Demyanenko, Andrew M. Giltrap, Benjamin G. Davis, Shabaz Mohammed

## Abstract

N-Hydroxysuccinimide (NHS) ester chemistry is used extensively across proteomics sample preparation. One of its increasingly prevalent applications is in isobaric reagent-based quantitation such as the iTRAQ (isobaric tags for relative and absolute quantitation) and TMT (tandem mass tag) approaches. In these methods, labelling on the primary amines of lysine residues and *N*-termini of tryptic peptides via amide formation (N-derivatives) from corresponding NHS ester reagents is the intended reactive outcome. However, the role of NHS esters as activated carboxyls can also drive the formation of serine-, tyrosine-, and threonine-derived esters (O-derivatives). These O-derivative peptides are typically classed as over-labelled and are disregarded for quantification, leading to loss of information and hence potential sensitivity. Their presence also unnecessarily increases sample complexity, which reduces the overall identification rates. One common approach for removing these unwanted labelling events has involved a quench with hydroxylamine. We show here that this approach is not fully efficient and can still leave substantial levels of unwanted over-labelled peptides. Through systematic screening of nucleophilic aminolysis reagents and reaction conditions, we have now developed a robust method to efficiently remove over-labelled peptides. The new method reduces the proportion of over-labelled peptides in the sample to less than 1% without affecting the labelling rate or introducing other modifications, leading to superior identification rates and quantitation precision.

## Introduction

NHS activated ester compounds were originally developed for peptide synthesis under diverse, including aqueous, conditions[1]. They are highly reactive carboxyl derivatives and when applied to proteins show some selectivity towards primary amines of *N*-termini and *N_Ɛ_* of lysines[2]–[4]; reversible reactions with disulfides and hydroxyl groups of tyrosines have been suggested[5]–[7]. NHS esters have therefore found widespread use in a range of biochemical applications including, for instance, covalent modifications for enzyme activation[5], [8], biotinylation for hybridisation and detection[9]–[11], heavy metal and fluorescent labelling for imaging and structure elucidation[2], [12].

In the proteomics field, NHS esters are used as cross-linking reagents for protein network analysis and structure elucidation[13]–[15] as well as to introduce stable isotope labels to tryptic peptides for quantitation[16], [17]. Specifically, tandem mass tags (TMT)[17] and isobaric tags for relative and absolute quantitation (iTRAQ)[16] have been developed to allow quantification of multiple samples in a single run, thereby minimising impact of technical variation. Both reagent types rely on NHS-ester chemistry to attach isotopically coded groups to *N*-termini and lysines of tryptic peptides of each sample before combining them for analysis. These modifications result in the same additional mass across all conditions but release a differential reporter ion upon collision induced dissociation (CID), which provides quantitative information. As the samples are mixed prior to analysis, complete labelling and robust quenching are paramount to precision of quantitation using this method.

The NHS ester labelling reaction at most *N*-termini and lysine ɛ-amines as reactive nucleophiles is considered to be most efficient at pH 8.0-9.0, under conditions where certain side products, such as tyrosyl esters, are formed and hydrolysed[18]. At this pH, the rate of concurrent NHS ester hydrolysis also increases and so is in competition with the desired amide labelling reaction. Together with slight differences in individual reaction kinetics due to the local environments of nucleophiles on peptides, small changes in reaction conditions can lead to incomplete and/or heterogeneous labelling[19]. As a result, peptide concentration, reaction pH, and water content have been optimised and under most current protocols achieve >99% labelling efficiency and minimise side product formation[20]. However, under these conditions, it has been shown that alongside tyrosines, serines and threonines can also be labelled though *O*-acylation of their hydroxyl groups to form ester O-derivative (scheme 1), especially when located in proximity to histidines[21]–[24] that may act as internal relay residues or general bases (Scheme 1). Together, these amino acids (Ser, Thr, Tyr, STY) comprise over 15% of the proteome and so are similar in abundance to primary amines in the sample. As *O*-acylation reaction is typically slower than *N*-acylation under the conditions required for complete *N*-acylation of primary amines, a lower proportion of these hydroxyl groups is thus labelled. This leads to generation of several forms of each STY-containing peptide, each with a different number of tags[23]. Moreover, where there are multiple hydroxyl residues on the same peptide, *O*-acylation at a single site can occur on different residues which may translate into differential chromatographic retention.

**Scheme 1.**
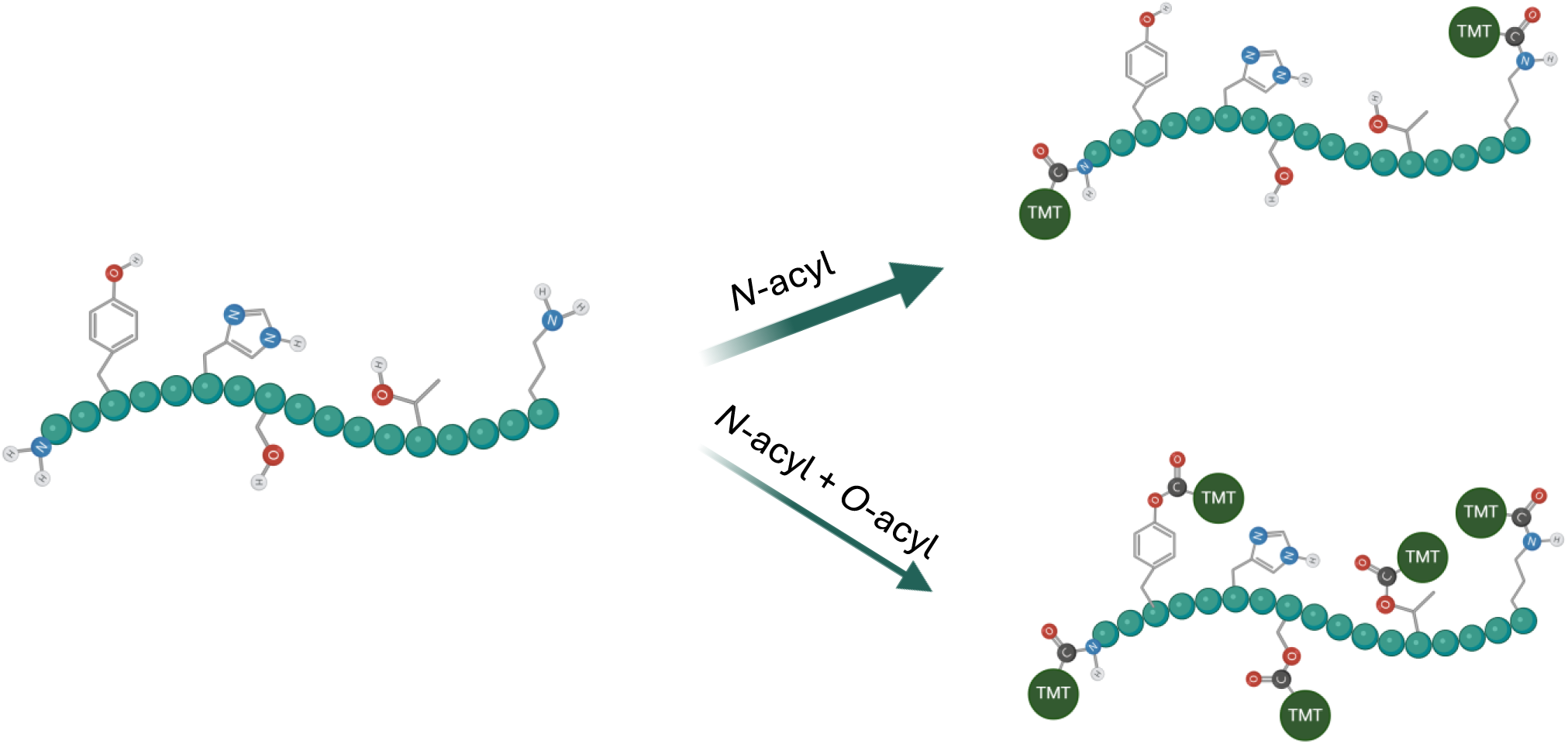
A schematic representation of potential products of peptide labelling with TMT reagent.

Regardless of origin, multiple versions of the same peptide unnecessarily increase sample complexity and lead to wasted time during data-dependent acquisition. Moreover, the stochastic nature of incomplete *O*-acylation in each sample may lead to increased variation and therefore decreased quantitation precision. These issues have been recognised by the community, and solutions have been sought. As *O*-acyl esters are less stable than the amide bonds generated through *N*-acylation of primary amines, it is potentially possible to selectively reverse *O*-acyl formation by employing an appropriate nucleophile. Hydroxylamine treatment was shown to reverse tyrosyl ester formation on model peptides and has since been universally applied to quench TMT labelling reactions[23]. However, like others before us, we have found that over-labelled peptides are still present in samples labelled and quenched with the standard protocol, and lead to systematic under-representation of STY-containing peptides in TMT datasets[20], [25]. Several alternative approaches have been proposed to further reduce the abundance of *O*-acyl esters in TMT-labelled samples. For example, raised temperature increases hydrolysis of *O*-acyl esters at high pH without introducing additional reagents. Thus, boiling of samples for 1 hour has been shown to reduce *O-*acylation in TMT-labelled samples[26]. Alternatively, others have proposed that there is reduced *O*-acyl ester formation when performing the labelling reaction at pH 6.7[25], [27]. However, under these conditions, the rate of *N*-acylation is also reduced, therefore complete labelling in solution requires higher temperature, excess of the labelling reagent and further reduced water content[25]. Therefore, whilst low pH labelling is able to drive complete labelling with TMT reagent-to-peptide w/w ratios of 1.6 : 1, this approach is not recommended for applied samples, where a higher ratio of 5:1 is suggested, making this method less economical[28]. As NHS ester compounds, including TMT reagents have been in use for decades, protocols for complete labelling of peptides have been optimized for many types of applications including single-cell protein quantitation[29] and automated workflows[30]–[33]. Required boiling of samples or change of pH between the digestion and labelling steps through desalting, drying and resuspension can, for example, introduce additional sample losses and variation and may not be viable for some applications.

Here, through the reconsideration of the fundamental deacylation chemistries that may be selectively applied to unwanted O-derivatives we have re-examined the strategy of cleavage post-labelling. The resulting discovery of the additional more efficient nucleophiles for this ‘clean-up’ of NHS-labelling reactions substantially reduces *O*-acylation with minimal protocol changes.

## Experimental Procedures

### Expi293F Lysate preparation and protein digestion

Expi293 (human cell line based on HEK 293) cell pellet was resuspended in lysis buffer (2% SDS (Sigma-Aldrich), 50 mM TEAB (Supelco) and sonicated using BioruptorPico (Diagenode) for 30 cycles (30s on/off). Protein concentration was determined with BCA assay (Thermo Fisher Scientific) prior to reduction and alkylation with 10 mM TCEP (Thermo Fisher) and 50 mM Chloroacetamide (Sigma-Aldrich) for 30 minutes. Proteins were then digested using a modified single-pot solid phase assisted sample preparation method (SP3)[34], [35] using magnetic carboxyl coated magnetic beads (Cytiva) at 4:1 w/w ratio. Proteins were precipitated onto the beads using acetonitrile (80% final concentration) and incubated for 30 min with shaking. Once proteins were immobilised, the beads were washed three times with 80% Ethanol, followed by three washes with 100% acetonitrile before incubation with digestion buffer (50 mM TEAB, containing Sequencing grade Trypsin (Promega) at 1:25 enzyme-to-protein for 4 hours. Supernatant was collected and beads were washed with 2% DMSO solution and combined with supernatant before acidification with formic acid (5% final concentration) and centrifugation at 16,000 x g for 10 minutes. Peptides were then desalted on Oasis HLB cartridges and eluted with 50% acetonitrile in ultrapure water. After desalting, the peptides were dried using Genevac EZ-2, and resuspended in 50 mM TEAB. The concentration of peptides was estimated using absorbance at 205 nm and adjusted to approximately 5 mg/ml before snap freezing in 100 ug aliquots for further use.

### TMT labelling and quenching

0.8 mg of TMTzero reagent (Thermo Fisher Scientific) was brought to RT and dissolved in 100 μl of dry acetonitrile under Ar gas. 20 μl of the TMTzero reagent in acetonitrile (160 mg TMT reagent, approx. 4-times molar excess to primary amines in this reaction) was mixed with the thawed 20 μl (100 μg) aliquot of tryptic peptides and incubated for two hours at room temperature with shaking. Where more peptide mixture was required, several labelling reactions were mixed and aliquoted for treatment. The reaction was quenched using different reagents and quantities specified for each experiment individually. Standard quenching condition was 1% final concentration of hydroxylamine (Sigma-Aldrich), equivalent to 0.3 M, for 30 minutes. For initial screening, final concentrations of *O*-Methoxylamine HCl (Sigma-Aldrich), Methylamine (in methanol, Sigma-Aldrich), Hydrazine hydrate (Sigma-Aldrich) were kept at 0.3 M. In methylamine treatment optimisation experiment, a 2-fold serial dilution of quenching reagent in methanol was made to result in 0.025 – 1.66 M final concentration, the reaction volume was set to 120 µL. The volume and solvent content of each reaction was kept constant by using 20 µL of labelled peptide mixture and adding 100 µL of quenching reagent solution in methanol. For evaluation of the final quenching protocol, three aliquots of 100 µg of peptides were labelled separately and split for quenching. 25 µg of TMT-labelled peptides were used for each reaction. Hydroxylamine and methylamine were used at 0.4 M, while 0.2 M Tris (Sigma-Aldrich), and 2% ammonium hydroxide (Fisher Scientific) were used to ensure robust pH control at 9.2 and above 11.5, respectively.

### LC-MSMS data acquisition

Peptides were separated on Ultimate 3000 RSLCnano system (Thermo Fisher Scientific) equipped with a C-18 PepMap100 trap column (300 μm ID x 5mm L, 100Å, Thermo Fisher Scientific) and an in-house packed Reprosil-Gold C-18 analytical column (50 μm ID x 500mm L, 1.9 μm particle size, Dr. Maisch). Mobile phases (A: 0.1% FA, 5% DMSO and 94.9% water; B: 0.1% FA, 5% DMSO, 94.9% ACN) were delivered at a flow rate of 100 nL/min. For single-shot LC-MS/MS analysis, 30, 60, or 120-minute gradient (10-36% or 10-32% B) was applied. Eluting peptides were electro-sprayed into an Orbitrap Ascend Tribrid mass spectrometer (Thermo Fisher Scientific), using Data-Dependent Acquisition mode (DDA).

For the initial screening experiments, full MS scans (350-1,400 m/z) were acquired in the Orbitrap analyser at 60,000 resolution with a 1.2 x 10^6^ AGC target, and 123 ms maximum injection time. Forty most intense precursors (charge states 2-7) from MS1 scans were selected and isolated at 1.2 Th with the quadrupole for MS/MS event using higher-energy collision dissociation (HCD) at a normalized collision energy (NCE) of 26%, and the fragmentation spectra were detected by ion trap using turbo scan rate mode (2 x 10^4^ AGC target and 32 ms maximum injection time). Dynamic exclusion of 20s was enabled. For method optimisation kinetic experiment and final method evaluation, the MS1 parameters were the same as above. MS2 scans were acquired in the Orbitrap at 7,500 resolution (4 x 10^4^ AGC target) and 64 ms maximum injection time). Dynamic exclusion of 25s was enabled.

### Database Searching

The raw files were searched against Uniprot Human database (Proteome ID: UP000005640) downloaded in August 2022 (79,759 sequences) using MSFragger search engine within FragPipe 21.1[36]. Three types of searches were conducted as appropriate: labelling efficiency search, over-labelling search, and off-target effects search. To evaluate labelling efficiency standard closed search parameters were used, with an additional TMTzero variable modification of 224.15248 Da on lysines and N-termini. To evaluate over-labelling, standard closed search settings were used with addition of the TMTzero label (224.15248 Da) as a fixed modification on N-termini and Lysine residues as well as a variable modification of the same mass on serines, threonines and tyrosines. MSBooster tool was enabled. To assess the extent of additional unpredicted modifications when using new quenching reagent, the default open search against the same database was used with the following modifications: TMT label was included as a fixed modification on lysine and peptide N-terminus, the range of MS1 delta masses was set to -225 to 300 to allow identification of under-labelled peptides[37]. IonQuant quantitation without signal normalisation was enabled for closed searches. PTM-shepherd (with default settings) was enabled for open searches to allow localisation of any discovered modifications[38].

### Experimental design and statistical rationale

For protocol development we sought to find trends in ester removal and focused on peptide population analysis in individual samples. Initial reagent screening experiments were conducted as three independent experiments using single replicates per condition. The rate of over-labelling in each sample was calculated based on sum of intensity of all over-labelled peptides. Representative results are shown. For method optimisation, 10 samples were made from a single peptide mixture labelled with TMTzero (Thermo Fisher Scientific): eight samples treated with a range of methylamine concentrations and two control samples. To record time-point data, aliquots of each sample were taken at specified times and acidified with formic acid (to 5% final concentration) to stop the reaction, generating a total of 50 samples for LC-MSMS analysis. The performance of the optimised protocol was evaluated using three separate TMTzero labelled peptide mixtures. Each labelled sample was split into 4 samples for quenching, generating a total of 12 samples. All samples were analysed using label-free single-shot LC-MSMS data-dependent acquisition runs. Where possible, over-labelling rates and labelling efficiency in each sample were calculated based on summed modified peptide intensities. To evaluate the effect of the quenching reagent on prevalence of over-labelled peptides and overall identification rates, a one-way ANOVA and Tukey’s pairwise comparison tests were used.

## Results

### Characterising the side reactions of NHS-ester-based labelling

We initially sought to evaluate the extent of side reactions, predominantly, *O*-acylation, that occur under typical NHS-ester mediated mass-tag labelling conditions. The most common use is in quantitative proteomic experiments, namely isobaric-based quantitation such as that obtained with TMT reagents. Therefore, we used a tryptic digest of a human cell line labelled with the TMTzero reagent as a model system [20]. As the quenching reagent concentration and reaction time vary across published labelling protocols (Supplementary table 1), we chose to quench with a conservative condition of 1% hydroxylamine and treatment time of 30 minutes, alongside no quenching. Samples were analysed using single-shot LC-MSMS runs. To evaluate the extent of *O*-acylation under these conditions, we calculated the proportion of peptide intensity associated with over-labelled peptides (Fig 1a). We found that after standard labelling without any quenching reagent, over 25% of peptides were over-labelled. In the sample treated with hydroxylamine, the proportion of over-labelled peptides was reduced, but only to 10%. The number of over-labelled peptides with more than one additional label was also substantially reduced in samples treated with hydroxylamine (Supplementary Figure S1). It is expected that serine, threonine and tyrosine will all be modified at different rates, and so we therefore compared the prevalence of *O*-acyl esters on each of these residues separately (Fig 1b). Without treatment, tyrosyl esters were the most prevalent of the three ester types, consistent with previous studies of *O*-acylation [23], [39]. While hydroxylamine treatment resulted in near complete removal of tyrosyl esters, it notably failed to substantially reduce the occurrence of serine and threonine esters. This is also in agreement with previously published data[20], [25], [40].

**Figure 1.**
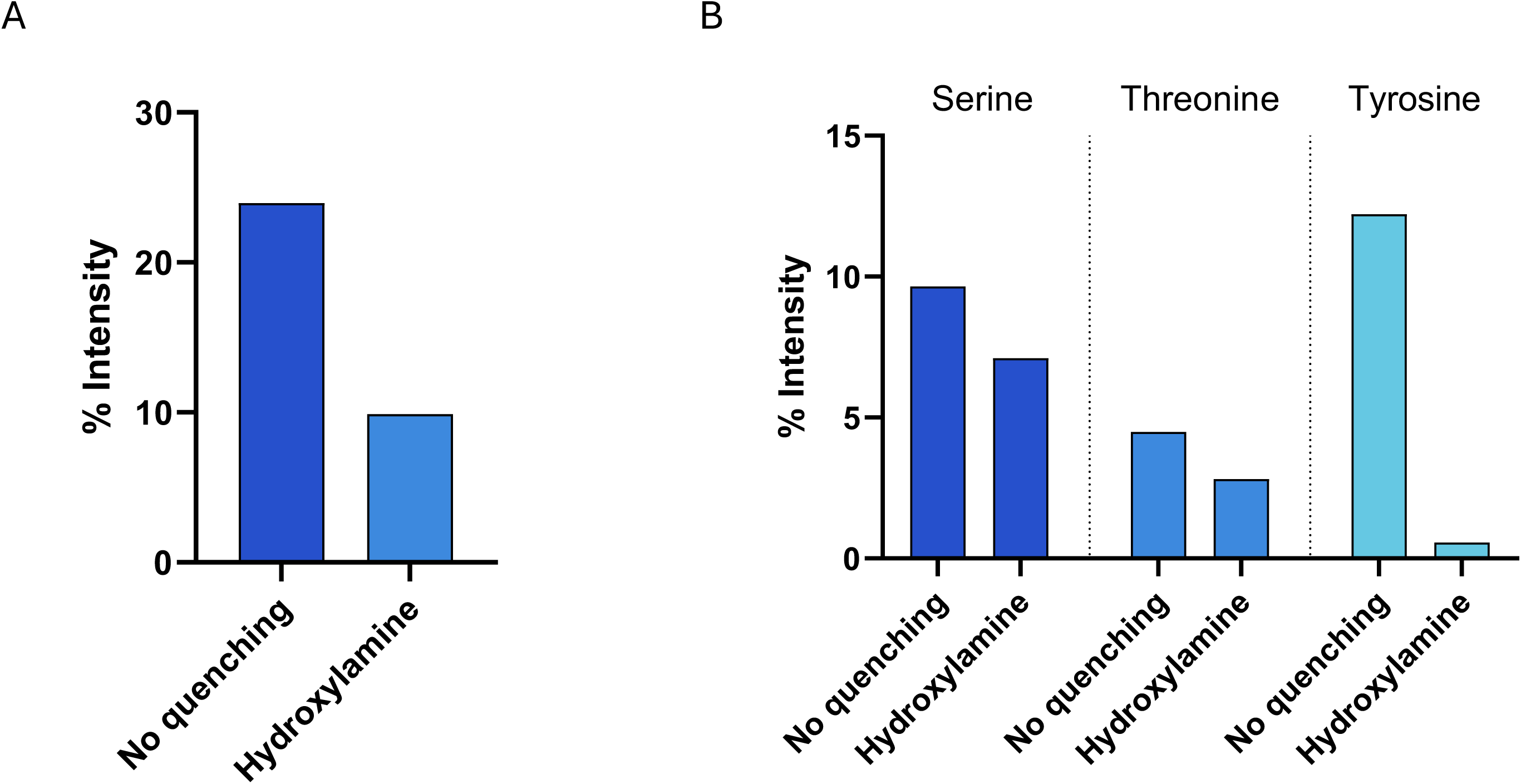
Evaluation of over-labelling using standard labelling and quenching protocols. (A) A bar chart of the percentage of peptide intensity from over-labelled peptides with and without standard hydroxylamine treatment. (B) Bar chart of the percentage of peptide intensity attributable to over-labelled peptides by ester type with and without hydroxylamine treatment

As the presence of histidines on peptides has been shown to increase the incidence of hydroxyl group reactivity towards NHS esters[20], [21], [24], [25], we also assessed whether the peptides over-labelled in our experiments contained histidine residues (Supplementary Figure S2). Consistent with previous studies, the incidence of over-labelling was higher in histidine-containing peptides from both treated and untreated samples (Supplementary Figure S2a). At the same time, despite histidine being the second least abundant amino acid in the proteome, histidine-containing peptides constituted a large proportion of all over-labelled peptides (Supplementary Figure S2b). Therefore, while hydroxylamine treatment reduced overall over-labelling, histidine containing peptides were disproportionately unaffected, suggesting that histidine not only plays a role in *O*-acylation but also in its maintenance under prior conditions used to remove it. Stable labelling of the histidine itself has been ruled out in the past[20], [21]; several reaction mechanisms have been proposed suggesting that the imidazole group of the histidine may instead play a catalytic role via the formation of transient acyl intermediates (nucleophilic catalysis) or via general base assistance [21], [24]. Indeed, similar approaches may be exploited to allow for selective modification of tyrosine residues at lower reaction pH, regardless of the peptide sequence by using excess of imidazole during labelling[22]. Analysis of sequence windows around over-labelled residues using probability logo plots[41], adjusted for natural probabilities in the background proteome shows enrichment of histidine residues particularly in +/-1-3 positions around each of the modified residues.

### Screen for superior clean-up reagent

When considering their deacylation, serine, threonine and tyrosine provide three different ester moieties and leaving groups that may therefore affect the rate of removal of the TMT label. The phenolate of tyrosyl aryl esters is the most labile. In contrast, serine and threonine derivatives, as alkyl esters, are more stable. Given the likely critical role of any putative nucleophile in not only determining the rate but also the nature of the rate-limiting or -determining step in deacylation we therefore sought to vary nucleophile. Only the esters of tyrosines were efficiently cleaved by hydroxylamine after 30 minutes of treatment.

Three reagents with varying nucleophilicity were used to ‘quench’ the TMTzero labelling reaction over varying time periods: *O*-methylhydroxylamine (methoxylamine), hydrazine hydrate, and methylamine[42]. To mimic the typical conditions that are used for hydroxylamine treatment, all reagents were applied at a concentration of 0.3M, and the reaction time was initially kept at 30 minutes (Fig 2a). The proportion of intensity associated with peptides containing additional labels on serine, threonine and tyrosine was used to estimate the extent of over-labelling. Methoxylamine had little effect, while quenching with hydrazine hydrate reduced over-labelling by 20% compared to the equivalent hydroxylamine treatment. Overall, methylamine was the most effective at removing *O*-acylation and halved total over-labelling to just over 5%.

**Figure 2.**
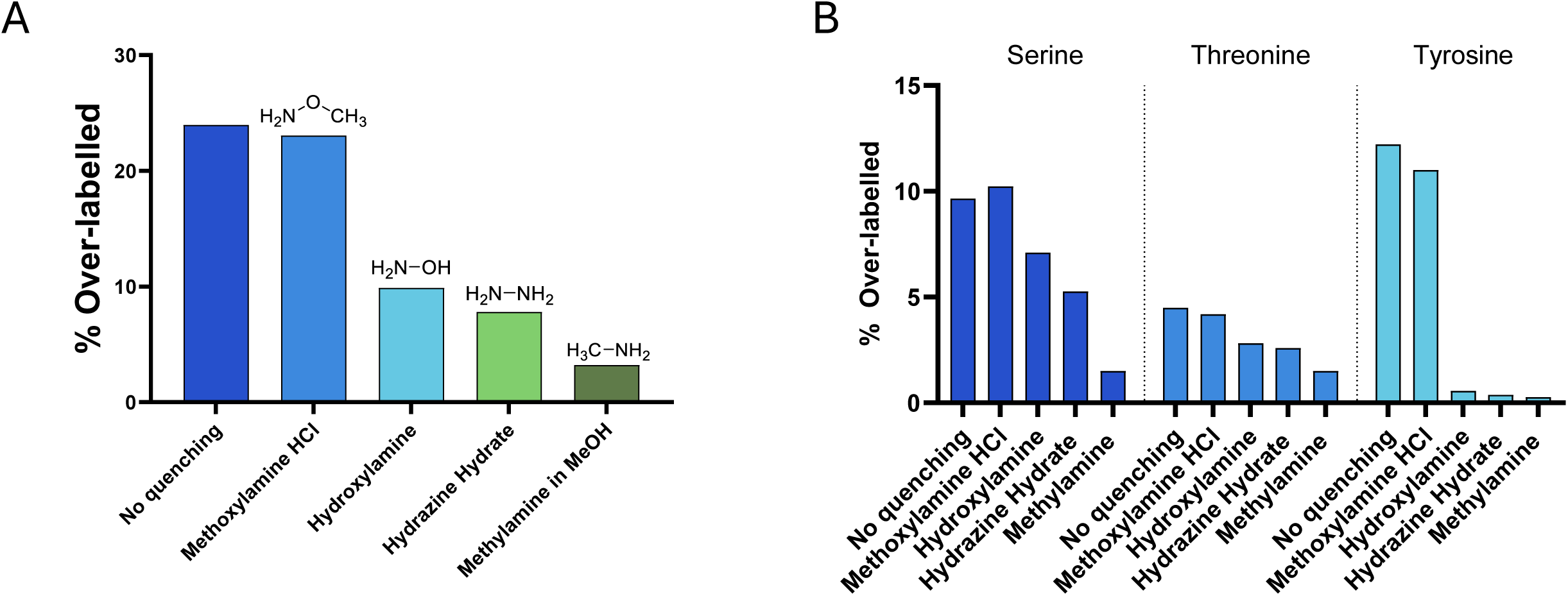
Screening of aminolysis reagents. (A) Bar chart showing the percentage of the intensity attributable to peptides containing at least one over-labelled residue after treatment with 50% acetonitrile (control), or 0.3 M of each: methoxylamine, hydroxylamine, hydrazine hydrate, and methylamine. (B) A bar chart showing the percentage of over-labelled peptide intensity by ester type after each treatment.

We evaluated the source of improvement in over-labelling by estimating initial rates for each of the three hydroxyl-containing amino acids separately (Fig 2b). While improvement was noted for both serine and threonine ester removal with more nucleophilic reagents, most of the improvement came from the reduction in *O*-acyl esters of serines, indicating that threonine esters are the most stable of the three ester types and require harsher conditions to be removed. This observation is consistent with the corresponding Taft parameters of substituted alkyl esters[43].

### Optimisation of methylamine clean-up conditions

As methylamine was the most successful reagent and showed promise in removing serine and threonine esters, we sought to achieve complete ester removal by optimising reaction conditions. We investigated the impact of reagent concentration and reaction time. We chose a range of methylamine concentrations from 0.013 to 1.66 M and monitored the reaction taking measurements at five timepoints: 15, 30, 60, 120, 240 minutes. Hydroxylamine and 50% acetonitrile were used as comparisons. For each treatment and timepoint, the proportions of total intensity assigned to correctly labelled peptides in each sample were calculated (Figure 3a). At the highest concentration of methylamine (1.66 M, approximately 2100 eq. to TMTzero reagent) we observed near complete removal of *O*-acyl esters after 15 minutes. Data suggested that at lower concentrations of methylamine, the same effect could be achieved by incubating the reaction for a proportionally longer time. At concentrations above 0.2 M, the abundance of correctly labelled peptides reached 99% in under 2 hours. In contrast, hydroxylamine treatment did not benefit from extended incubation time and the proportion of correctly labelled peptides plateaued at 92% after 1 hour, further highlighting the need for an alternative reagent.

**Figure 3.**
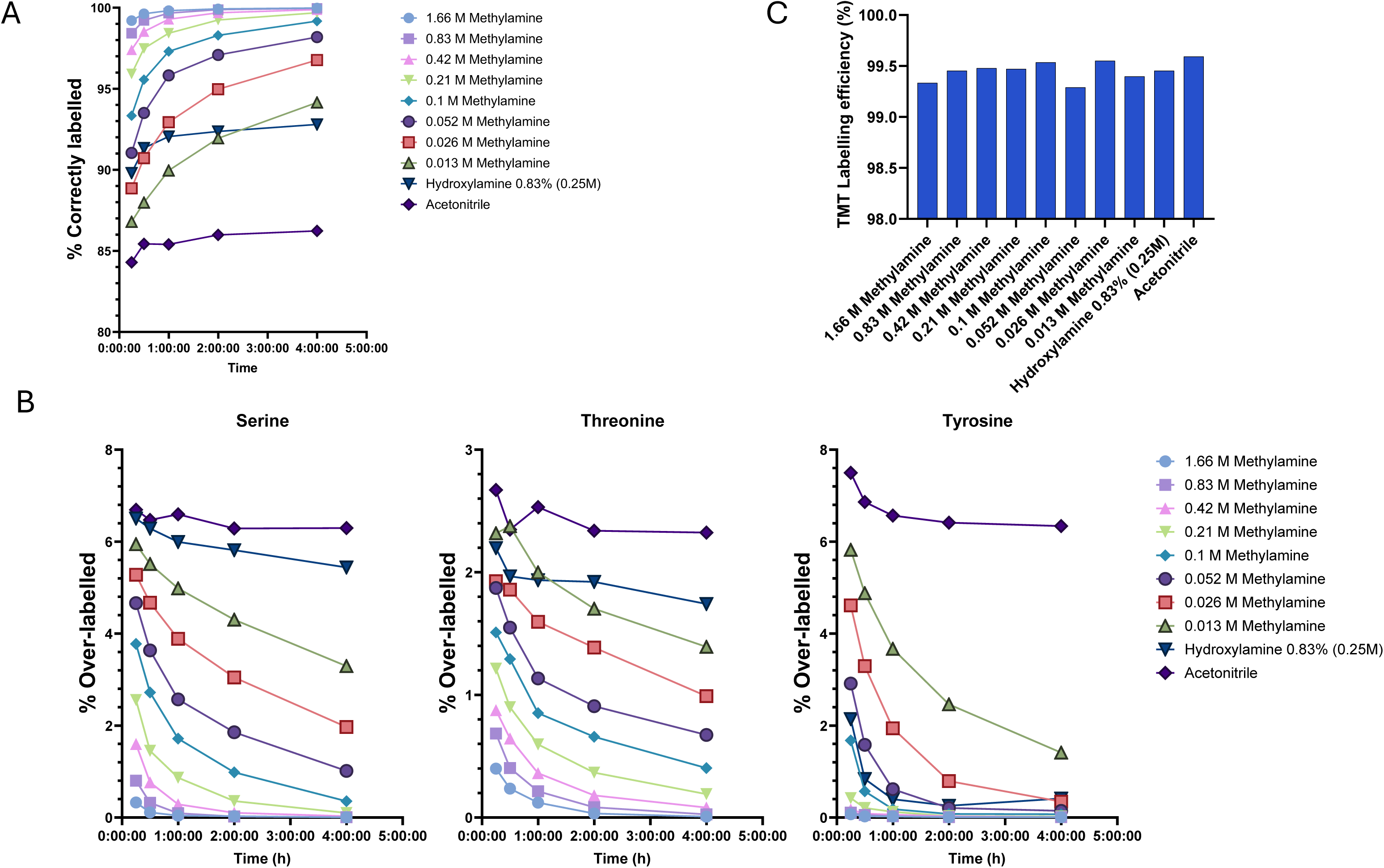
Optimisation of methylamine concentration and reaction time to minimise the proportion of over-labelled peptides. (A) A line graph showing the proportion of peptides (by intensity) correctly labelled only at lysines and N-termini after 15 min to 4 hours of quenching with increasing concentrations of methylamine compared to standard treatment with hydroxylamine and no quenching controls. (B) A line graph showing the proportion of peptides (by intensity) containing *O*-acyl esters of serines, threonines and tyrosines for each treatment and timepoint. (C) A bar chart showing the proportion of labelled lysines and N-termini (labelling efficiency) after 2 hours of treatment.

To confirm that the improvement in over-labelling in methylamine treated samples was due to additional removal of serine and threonine esters, we separately plotted the prevalence of the modification on each of the residues (Figure 3b). As expected, both standard hydroxylamine treatment, and methylamine treatment at concentrations over 50 mM were effective at removing tyrosyl esters in under 2 hours. Albeit methylamine was over four times as efficient as hydroxylamine, requiring four-to-five times lower concentration or incubation time to achieve the same result. The improvement in removal efficiency of seryl-and threonyl-esters was even more striking. Whereas hydroxylamine failed to substantially reduce the abundance of these esters even after 4 hours, a similar concentration of methylamine more than halved the abundance of serine and threonine-esters after 30 minutes; once again, threonine esters appear to be the most resistant of the three ester types. At methylamine concentrations above 0.4 M, relative abundance of remaining over-labelled peptides was less than 0.25% after 2 hours of treatment.

Considering the higher basicity of methylamine and the resulting increase in sample pH from ∼8.5 to above 11, the possibility of new unforeseen reactions developing over the extended incubation time was also considered. Cleavage of amide bonds at lysines and peptide *N*-termini generated through NHS labelling with TMT reagent is highly unlikely, but nonetheless possible. To exclude this, we searched the 2-hour timepoint data using the TMT label as a variable modification and calculated the labelling efficiency. We found that over 99% of all identified peptides were labelled at lysines and *N*-termini regardless of treatment, indicating that the amide bonds created during the labelling reaction remain stable (Figure 3c).

To rule out the formation of other unexpected products, we used the open search functionality of MSFragger to detect any new modifications on peptides after two hours of treatment[37]. Although we identified over 300 potential mass shifts, the majority of these were represented by 0-10 peptide spectrum matches (PSMs) per sample and were also present in the control experiments (Supplementary table S2). To assess whether methylamine had any concentration-dependent effect on the prevalence of unexpected modifications, we filtered the delta mass list to only those with more than 10 PSMs in the sample treated with 1.66 M methylamine and calculated the difference in total percentage of PSMs in each sample compared to the untreated control (Supplementary Figure S3). Overall, the prevalence of PSMs with additional delta masses decreased with increasing concentrations of methylamine. This was largely due to the reduction in over-labelling caused by methylamine treatment from over 8% of all PSMs in untreated sample to just over 0.1% in the sample treated with 1.66 M methylamine. The abundance of other prominent delta masses including under-labelling, deamidation and methylation at asparagines, and a delta mass of –198.136 Da with unknown composition only showed small differences between conditions (Supplementary Figure S3). The best overall decrease in side-products of the TMT labelling reaction after 2 hours of quenching was in samples treated with 0.2-0.8 M methylamine. To minimise treatment time, 1 hour incubation and 0.4M methylamine was used in further experiments.

### Performance of optimised quenching protocol

We next sought to confirm the robustness of the new protocol and evaluate the extent of its benefit on peptide identification rates. We labelled 100 μg aliquots of peptides with TMTzero reagent and quenched the reaction with 0.4 M of either hydroxylamine or methylamine for 1 hour (Figure 4a). As the final pH of the reaction is influenced by the quenching reagent, 0.2 M Tris pH 9.2 and 2% ammonium hydroxide (approx. final pH 11.6) buffer solutions were used as untreated controls to rule out any effect due to alkaline hydrolysis. The extent of over-labelling was assessed by comparing the sum of intensities of the peptides modified on serines, threonines, or tyrosines to the total signal (Figure 4b). Although the pH change alone had a small effect on the abundance of over-labelled peptides, as expected, both hydroxylamine and methylamine treatments resulted in significantly less over-labelling than respective controls. Methylamine was significantly more effective than hydroxylamine with less than 1% *O*-acylated peptides (p-value < 0.0001). The labelling efficiency of primary amines was not affected by any of the treatments and after one hour was >99% in all samples (Supplementary Figure S4).

**Figure 4.**
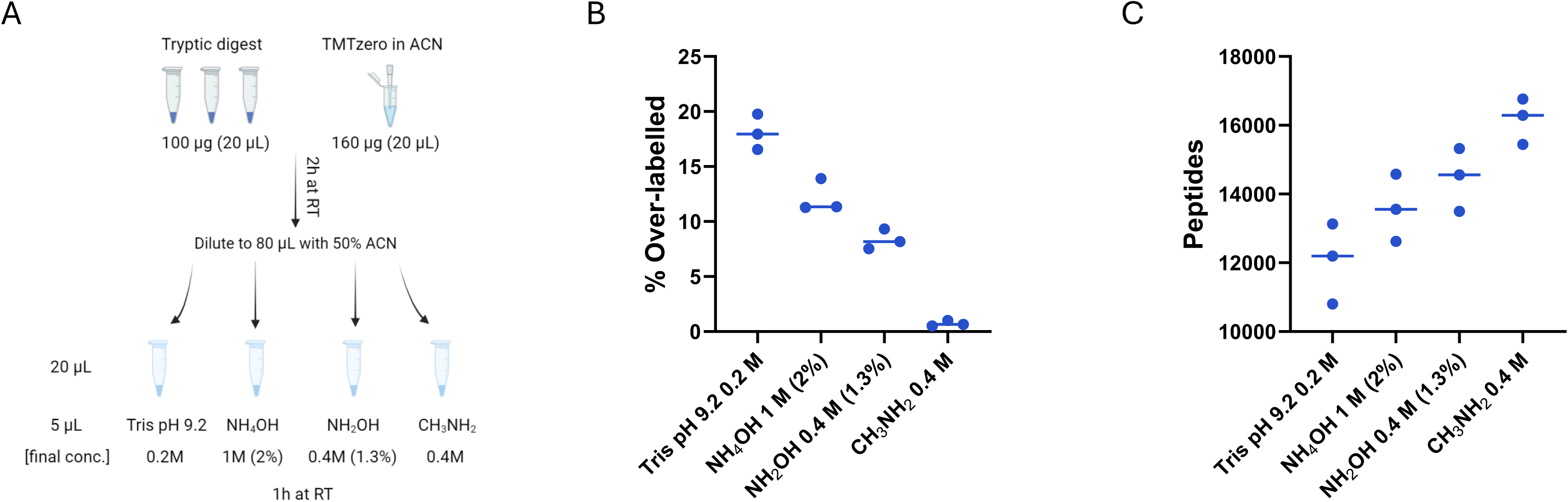
Evaluation of the optimised quenching protocol. (A) Schematic representation of experimental design. Peptides were labelled with standard protocol (samples and reagent amounts and volumes are indicated). Tris, ammonium hydroxide and hydroxylamine were used for comparison to optimised methylamine method, final reagent concentrations, volumes, and reaction time are indicated. (B) A graph showing the proportion of peptides containing O-acyl esters after each treatment. Individual values represent replicates, line represents the median. (C) A graph showing the number of peptides identified in single-shot LC-MS runs for each sample. Each point represents individual replicates, and lines are drawn at the median value

We hypothesised that this reduction in over-labelling would lead to an increase in peptide identification numbers in a standard TMT experiment due to the reduction in sample complexity. To evaluate the effect of the new quenching protocol on peptide identification, we searched the files using the same settings as those normally used for quantitative TMT experiments, i.e. treating labelling at peptide *N*-terminus and lysines as fixed modifications (Figure 4c). As expected, the average number of peptide identifications increased with hydroxylamine treatment, and methylamine treatment compared to the controls quenched with Tris or ammonium hydroxide. Methylamine treatment significantly increased the number of peptide identifications by up to 30% compared to the Tris-quenched control and provided an additional 10% improvement compared to hydroxylamine treatment.

## Discussion

Although widely regarded as amine-specific reagents, NHS esters have been shown to form significant amounts of *O*-acyl esters within peptide and protein mixtures. We have shown here that an alternative nucleophilic reagent, methylamine, can efficiently cleave all three types of *O*-acyl esters in a concentration-dependent manner (Figure 3b). Under optimised conditions, near complete removal can be achieved regardless of the initial over-labelling rate (Figure 4b). Focusing on selective removal of by-products post-labelling allows users to maintain previously developed robust labelling protocols with minimal changes to sample manipulation. The effect of such a clean-up is therefore two-fold for data acquisition and analysis: i) reduction in variation between correctly labelled peptides and ii) the overall reduction in sample complexity through signal consolidation. We observed the reduction in variation when comparing initial over-labelling rates (in Tris-quenched samples) and those treated with the optimised quenching protocol (Figure 4b).

The reduction in complexity through ester removal allows more instrument time for sequencing new peptides as evidenced by the increase in identification rates of correctly labelled peptides coinciding with the reduction in over-labelling (Figure 4b, c). Thus, this method is especially useful for challenging TMT-based workflows such as high-throughput applications involving heterogenous samples, sample-limited studies, and experimental designs involving multiple mixtures that rely on homogenous labelling within batches that can be compared to each other through a reference channel. Peptide-level quantitation will benefit more than protein-level quantitation experiments. For example, TMT-based phospho-proteomics workflows would particularly benefit since quantitation is performed at the site level based on peptide level data. Other areas within proteomics that utilise NHS-esters may also potentially benefit, for example, in cross-linking experiments relying on NHS esters. Inducing the formation of linkages involving STY residues and subsequent selective cleavage of tyrosyl linkages at low pH, followed by serine and threonine linkage cleavage by methylamine may help improve overall cross-linked peptide identification and site localisation.

In summary, we demonstrate that the simple replacement of hydroxylamine with methylamine can substantially improve ‘clean-up’ of NHS-ester mediated mass-tag labelling experiments and leads to significant benefits for the subsequent mass spectrometric data acquisition, as shown for TMT-based experiments. We anticipate that this minor modification, which is readily employable, will lead to considerably superior data for a substantial number of proteomics experiments.

## Abbreviations

AGC: Automatic gain control
BCA: Bicinchoninic acid protein concentration assay
CID / HCD: Collision induced dissociation / higher energy
CID DDA: Data-Dependent Acquisition
DMSO: Dimethyl sulfoxide
iTRAQ: isobaric tags for relative and absolute quantitation
MSMS / LC-MSMS: Tandem mass spectrometry, liquid chromatography coupled to MSMS
NCE: Normalised collision energy
NHS: *N*-Hydroxysuccinimide
PSM: Peptide spectrum match
SDS: Sodium dodecyl sulphate
SP3: Single pot solid-phase assisted sample preparation
TCEP: tris(2-carboxyethyl) phosphine
TEAB: Triethylammonium bicarbonate
TMT: tandem mass tag

## Acknowledgments

Chemistry (S.M, A.M.G, Y.D, and B.G.D.) at the Rosalind Franklin Institute is supported by the EPSRC (V011359/1 (P)).

## Author contributions

Y. D, S. M, and A. M. G designed experiments, Y. D performed experiments, Y. D analysed data, Y. D prepared figures, Y. D, S. M, A. M. G and B. G. D wrote the paper.

## Data availability

All raw data and search results presented in this paper have been deposited to ProteomeXchange Consortium via the PRIDE partner depository with the dataset identifier PXD054730.

## Supplemental data

This article contains supplemental figures and tables

## Supplementary tables

**Table S1.**
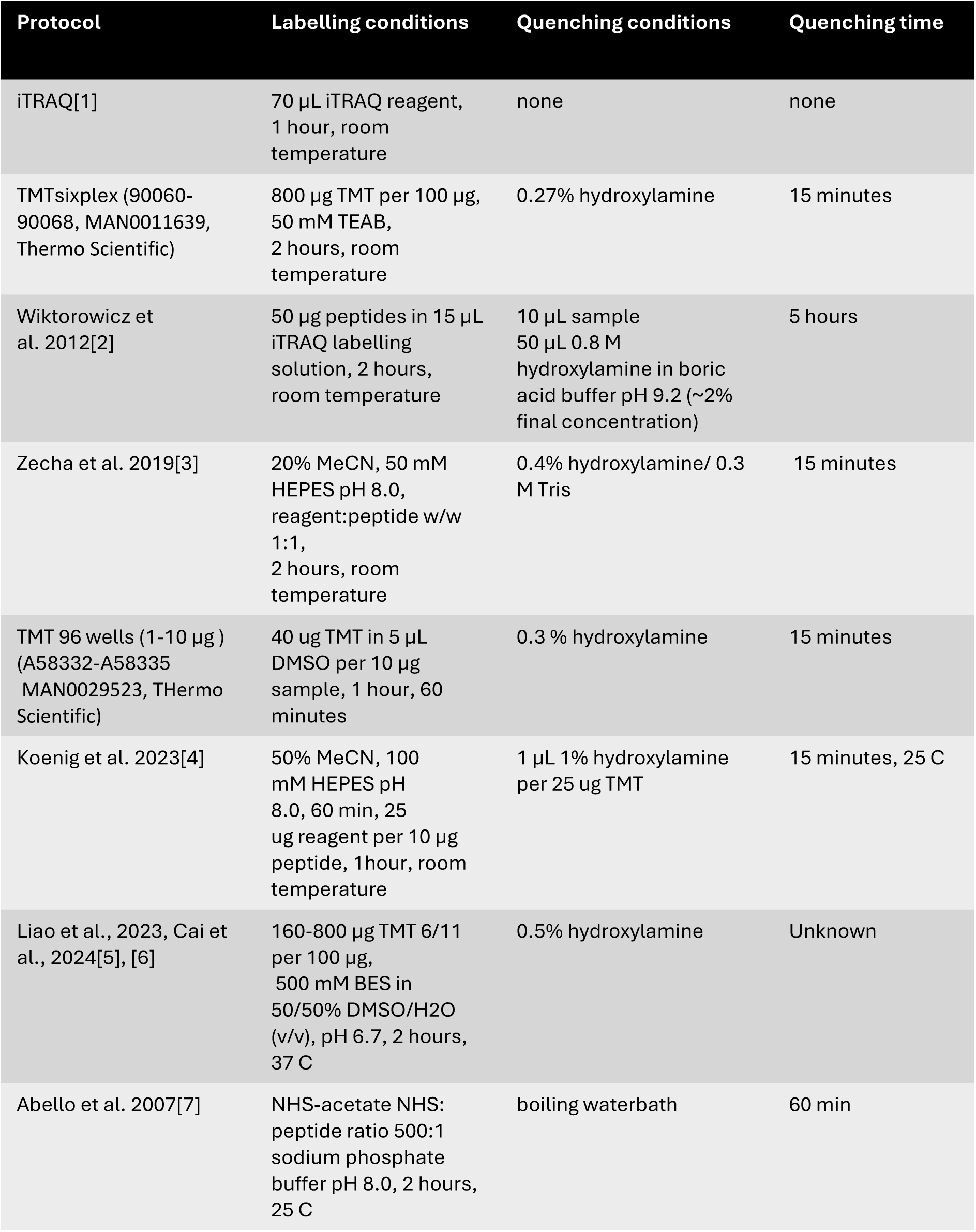
Overview of published isobaric labelling and quenching conditions.

**Table S2.** Table of all modifications identified in an MSFrager[8] open search, summarized with PTM-Shepherd[9].

**Figure S1.**
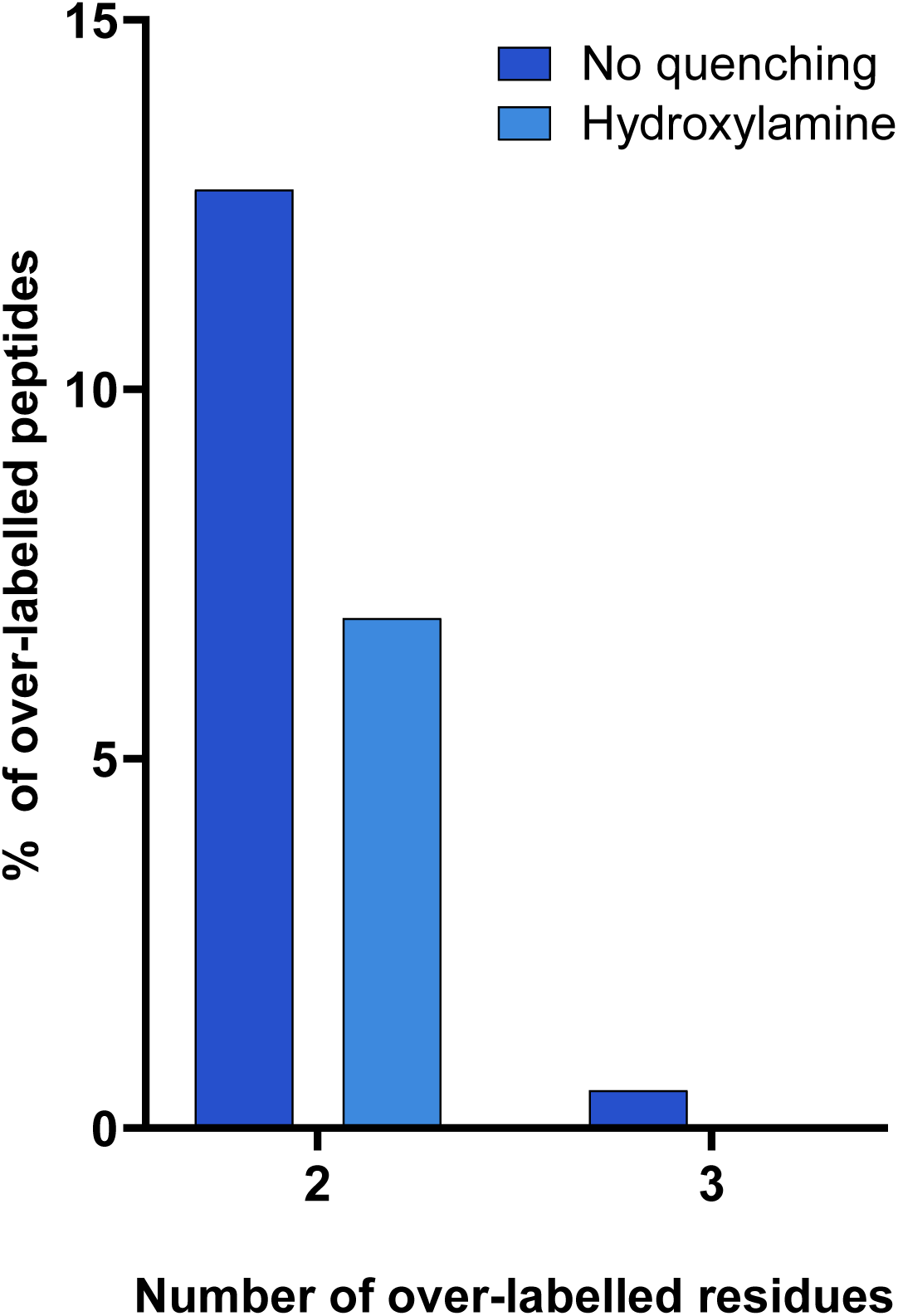
Analysis of over-labelled peptides by number of additional TMT-labels. A bar chart showing the percentage of over-labelled peptides containing more than one additional modification on serines, threonines or tyrosines with and without hydroxylamine treatment.

**Figure S2.**
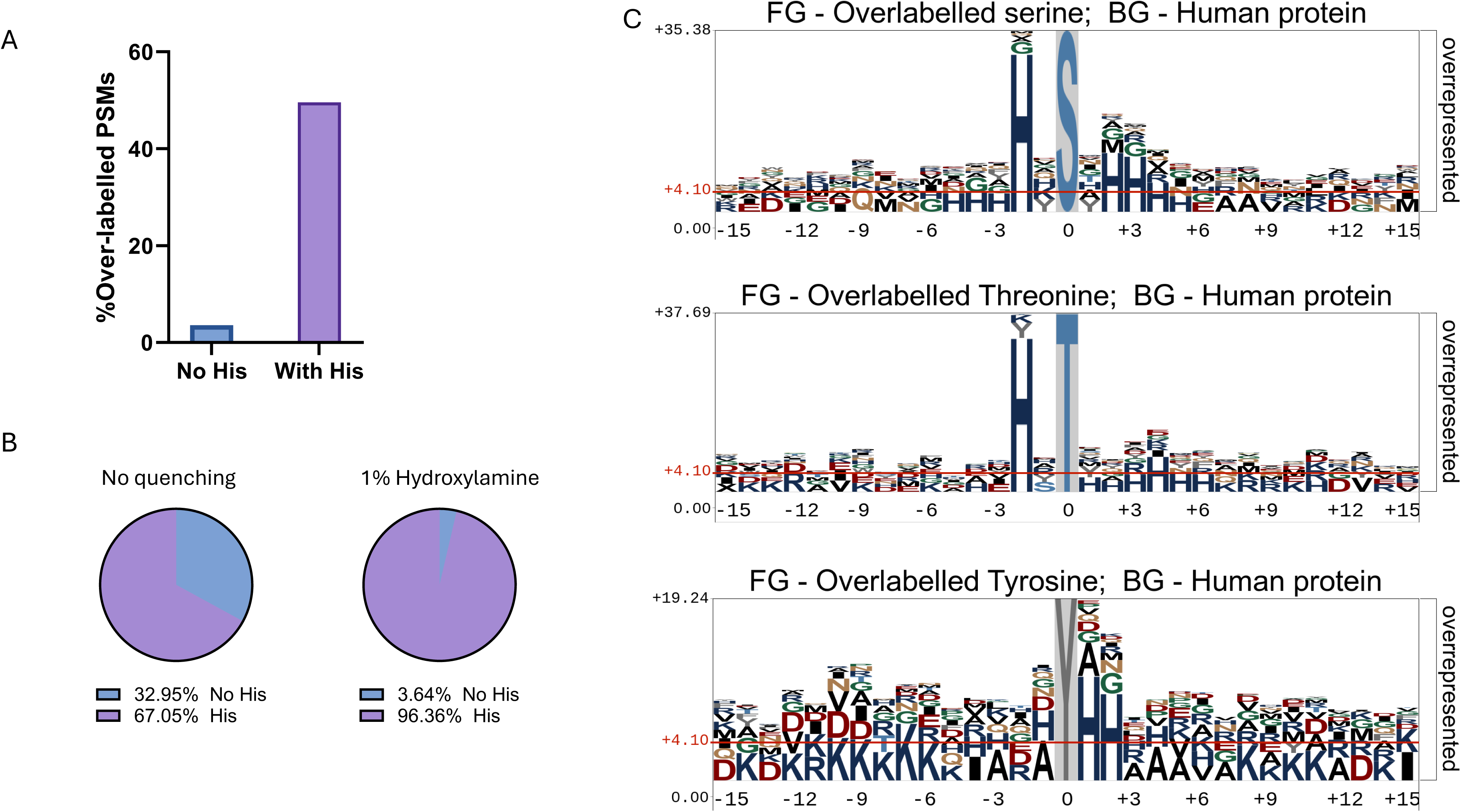
Sequence analysis of over-labelled peptides. (A) A bar chart showing percentage of spectrum matches to over-labelled peptides for sequences with and without histidine. (B) A pie chart showing the percentage of peptide sequences containing histidine in over-labelled peptides identified in untreated and hydroxylamine-quenched samples. (C) pLogo plots[10] of over-labelled peptide sequence windows separated by ester types and adjusted for natural residue occurrence in human proteome. Bottom row letters larger than the significance threshold are deemed significantly enriched in those positions.

**Figure S3.**
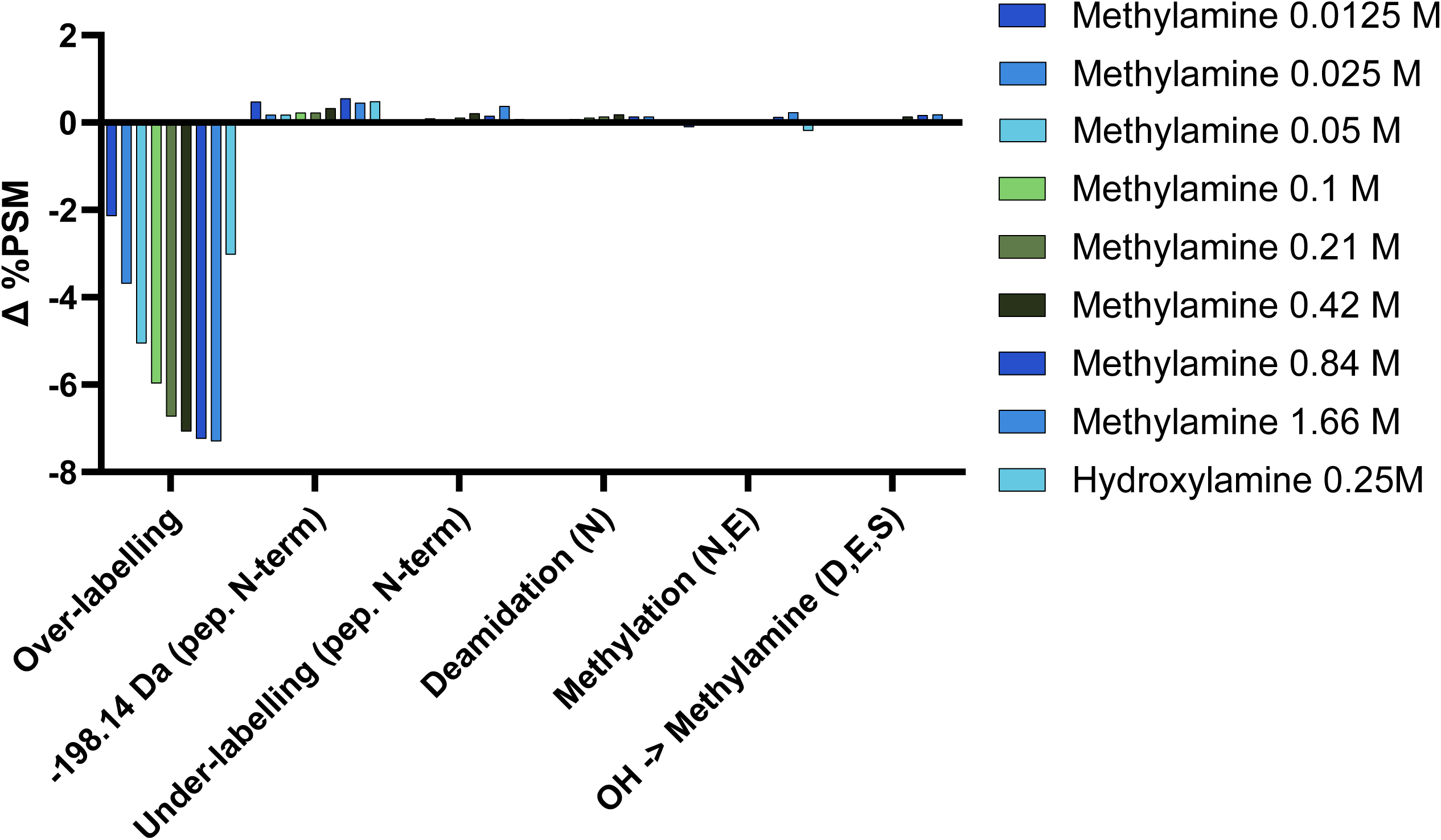
MSFragger open search results[8]. A bar chart of the difference in the percentage of peptide spectrum matches identified in methylamine (0.013 - 1.66 M) and hydroxylamine treated samples compared to the untreated control.

**Figure S4.**
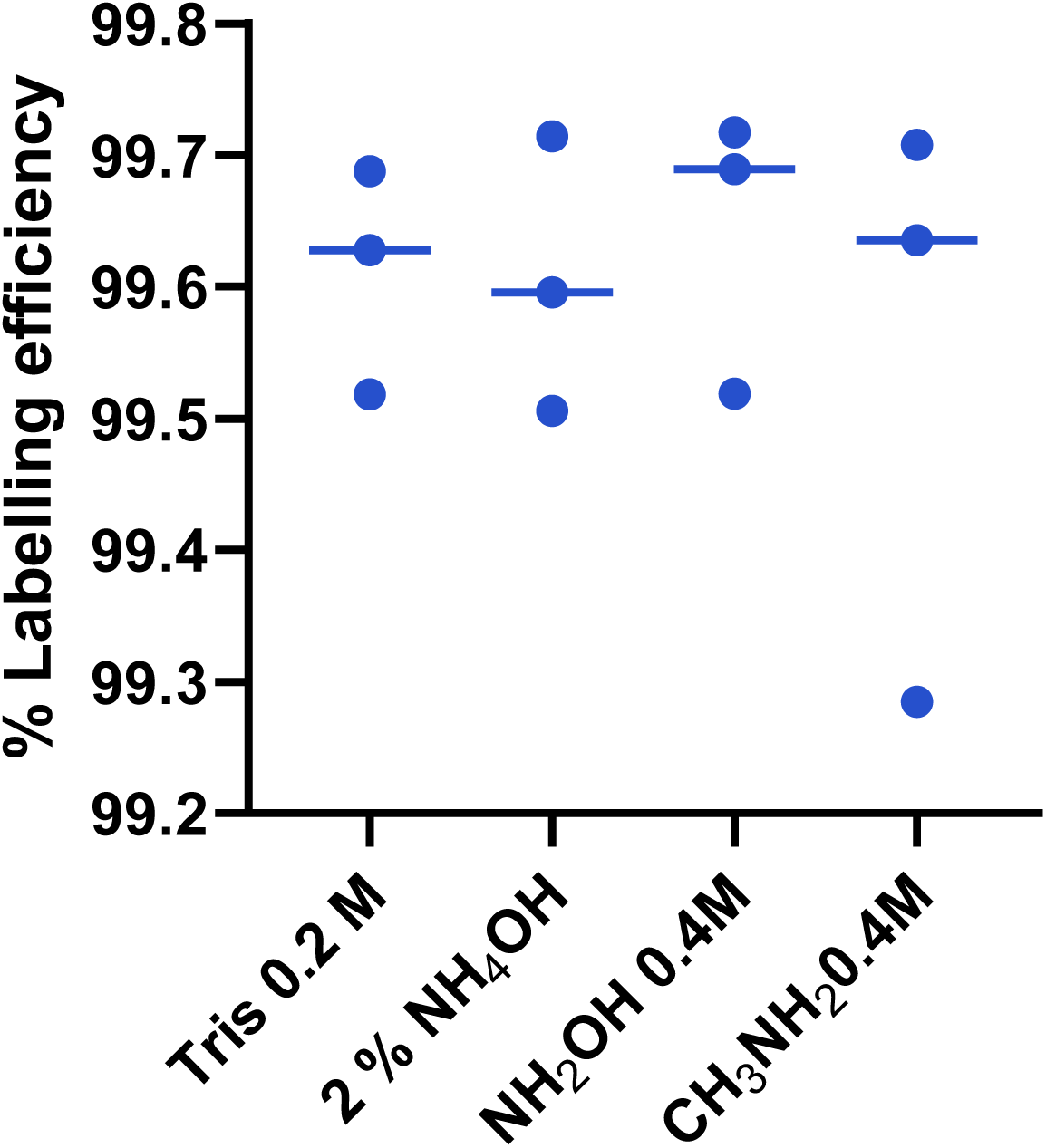
TMT-labelling efficiency in samples quenched for 1 hour with Tris (pH 9.2), Ammonium hydroxide (2%, pH ∼ 11.6), hydroxylamine (0.4 M, pH ∼8.5), methylamine (0.4 M, pH ∼ 11.5). Each point represents an individual replicate, with a line going through the median.

